# Isotope geolocation and population genomics in *Vanessa cardui:* Short- and long-distance migrants are genetically undifferentiated

**DOI:** 10.1101/2023.12.10.569105

**Authors:** Megan S. Reich, Daria Shipilina, Venkat Talla, Farid Bahleman, Khadim Kébé, Johanna L. Berger, Niclas Backström, Gerard Talavera, Clément P. Bataille

## Abstract

The painted lady butterfly *Vanessa cardui* is renowned for its virtually cosmopolitan distribution and the remarkable long-distance migrations that are part of its annual, multi-generational migratory cycle. Recently, *V. cardui* individuals were found north and south of the Sahara in the autumn, suggesting distinct migratory behaviours within the species. However, the evolutionary and ecological factors shaping these differences in migratory behaviour remain largely unexplored. Here, we performed whole-genome resequencing and analysed the hydrogen and strontium isotopes of 40 *V. cardui* individuals simultaneously collected in the autumn from regions both north and south of the Sahara. Our investigation revealed two main migratory groups: (i) short-distance migrants, journeying from temperate Europe to the circum-Mediterranean region and (ii) long-distance migrants, originating from Europe, crossing the Mediterranean Sea and Sahara, and reaching West Africa, covering up to over 4,000 km. Despite these stark differences in migration distance, a genome-wide analysis revealed that both short- and long-distance migrants belong to a single intercontinental panmictic population extending from northern Europe to sub-Saharan Africa. Contrary to common biogeographic patterns, the Sahara is not a catalyst for population structuring in this species. No significant genetic differentiation or signs of adaptation and selection were observed between the two migratory phenotypes (pairwise F_ST_ = 0.001 ± 0.006). Nonetheless, two individuals, which were early arrivals to West Africa and covered longer migration distances, exhibited some genetic differentiation. The lack of genetic structure between short- and long-distance migrants suggests that migration distance in *V. cardui* is a plastic response to environmental conditions.

**Significance statement:** Although migratory insects dominate living biomass fluxes and impact agriculture, ecosystems, and human communities, little is known about the controls of their migratory behavior. Our study develops an interdisciplinary framework, applied to the migratory butterfly *V.cardui*, to explore the genetic basis of variation in insect migration behavior. We leverage new generation isotope geolocation techniques to uncover striking differences in butterfly behaviour, with some individuals migrating short distances within the circum-Mediterranean region and others migrating thousands of kilometers across the Mediterranean Sea and Sahara. This major difference does not coincide with genetic differentiation or population structure and is likely a plastic response to environmental cues. This study provides a ground-breaking framework to study migration in insects.

## Introduction

Migration has evolved several times in insects (1) and is often viewed as an advantageous adaptation, as insects move to favourable environmental conditions, exploit seasonal resources, and escape parasites and predators (2). Many migratory animals show variation in migratory behaviour that results in distinctive spatiotemporal movement patterns. As an innate behaviour in insects, the proximate mechanisms for migration are likely held in the genome (3,4), but the environmental and genetic factors driving intraspecific variation in insect migratory behaviour are only beginning to be understood (5).

The painted lady butterfly *Vanessa cardui* is renowned for its obligate, annual, multi-generational migratory cycles throughout Africa, Asia, Europe, and North America (6,7). As part of a migratory cycle covering between 8 and 10 generations and involving both the Palearctic and Afrotropical regions, late summer and autumn generations migrate southward from temperate Europe towards more suitable breeding grounds. While a substantial portion of these southward migrating butterflies journey across the Mediterranean Sea and Sahara to subsequently breed in sub-Saharan Africa (8–10), others migrate to the circum-Mediterranean region (11,12). The factors driving these differences in migratory behaviour, resulting in the disjunct winter distribution of *V. cardui* across the Sahara, remains unknown.

Migratory phenotypes include observable characteristics such as migratory propensity, timing, orientation, and wing morphology (13). However, migratory behaviours are complex traits and are therefore difficult to quantify, especially in insects. The distance an individual butterfly covers during migration is a consequence of this complex phenotype, which creates a distribution of migration distances across individuals, reflecting the extent of the species’ ecological niche. In cases where the suitable niche is geographically continuous, it is expected that individuals at the extremes of the migration distance distribution (i.e., those who migrate the shortest or longest possible distances) are selected against due to the inherent risks associated with extremes. In such scenarios, stabilising selection may favour migrants that cover intermediate distances. Conversely, when the suitable niche is fragmented by large biogeographic barriers, such as the Sahara, extreme migration distances may be favoured by disruptive selection. Selection for specific migration distances can have important repercussions for survival and growth. For example, pronounced climatic differences between regions north and south of the Sahara influence the composition and quantity of host plant species, predators, parasitoids, and weather events, as well as influencing development time (9). Altogether, this scenario leads us to ask: can we detect the action of adaptive evolutionary forces acting on the extremes of the migration distance distribution and effectively disentangle their action from neutral evolutionary processes?

Differences in migratory routes, distances, and orientation has led to genetic differentiation and subsequent divergent selection in multiple bird species, such as the European blackcap *Sylvia atricapilla,* Swainson’s thrush *Catharus ustulatus,* and willow warbler *Phylloscopus trochilus* (14–17). Likewise, differences in migration distance could lead to similar spatiotemporal separation of *V. cardui* lineages, resulting in reduced genetic exchange between *V. cardui* that migrate across the Sahara and those that remain in the circum-Mediterranean region, thus leading to genetic differentiation. However, exploring the genetic underpinnings of natural variation in migratory behaviour requires not only whole-genome sequencing data, but also accurate migratory phenotyping (4). Studies on insect migration have lagged behind those of birds and mammals because identifying migratory phenotypes in insects is challenging (18). Extrinsic markers such as biologgers commonly record migratory trajectories for birds and mammals, but few studies have used these techniques on insects (19-21) because insects are generally too small, short-lived, and numerous for these techniques to be applied on a large scale. Instead, studies on insects have mainly relied on observations of physiological state (e.g., wing wear scores), behaviour (e.g., insect-monitoring radar), or regional field monitoring (e.g., spatiotemporal location).

Fortunately, naturally-occurring isotopes are intrinsic markers that have proven increasingly useful for estimating the geographic origin of wild-caught insect migrants and have the potential to identify migratory phenotypes. Isotope geolocation relies on the observation that insect larvae ingest local environmental isotopic signatures through their diet and incorporate these isotopic signatures into their developing tissues (22,23). Insect wings are relatively inactive tissues and largely preserve the isotopic signature of the natal environment (24,25). Thus, the isotopic signature of the wing of a wild-caught insect can be measured and then compared to a spatial model of isotopic variation (i.e., an isoscape) to estimate the insect’s natal origin, a process known as geographic assignment. However, single isotope-based geographic assignment generally yields broad and unspecific regions of natal origin making it challenging to calculate downstream estimates, such as migration distance. More spatially-constrained estimates of natal origin can be obtained by combining multiple isotopes with complementary patterns (26). Dual isotope-based geographic assignment combining hydrogen isotope values (δ²H), driven by climatic variables (e.g., precipitation; 23,27), and strontium isotope ratios (^87^Sr/^86^Sr), primarily driven by geological processes (e.g., bedrock type and age; 28), have previously been used to classify migratory behaviours in birds and bats (e.g., 29,30). However, isotopes have not often been leveraged to phenotype migratory behaviour and investigate the potential genomic drivers of variation in insect migratory behaviour.

Here, we combined isotope-based phenotyping of migration distance with whole-genome resequencing data to investigate the mechanisms driving variation in migratory behaviour for *V. cardui* individuals. By coupling δ²H and ^87^Sr/^86^Sr into a dual geographic assignment framework, we estimated the natal origin and migration distance of 40 adult *V. cardui* synchronically collected from north and south of the Sahara in the autumn. A genome-wide population resequencing approach was used to investigate potential genetic structuring associated with contrasting migratory phenotypes. This study endeavours to shed light on the underlying factors shaping the remarkable trans-Saharan migratory journeys of *V. cardui*.

## Results

### Isotope-based phenotyping reveals differences in migration distance

We estimated the natal origins of 40 *V. cardui* of unknown origin captured during the autumn southward migration from both north (i.e., Morocco, Spain, Portugal, Malta) and south (i.e., Benin, Senegal) of the Sahara using δ²H and ^87^Sr/^86^Sr following the geographic assignment framework of Reich *et al.* (26) (S1 Table; Fig 1D). From the posterior probability surfaces, we then calculated a conservative estimate of migration distance (i.e., minimum distance; S1 Fig; Fig 1C).

**Fig 1.**
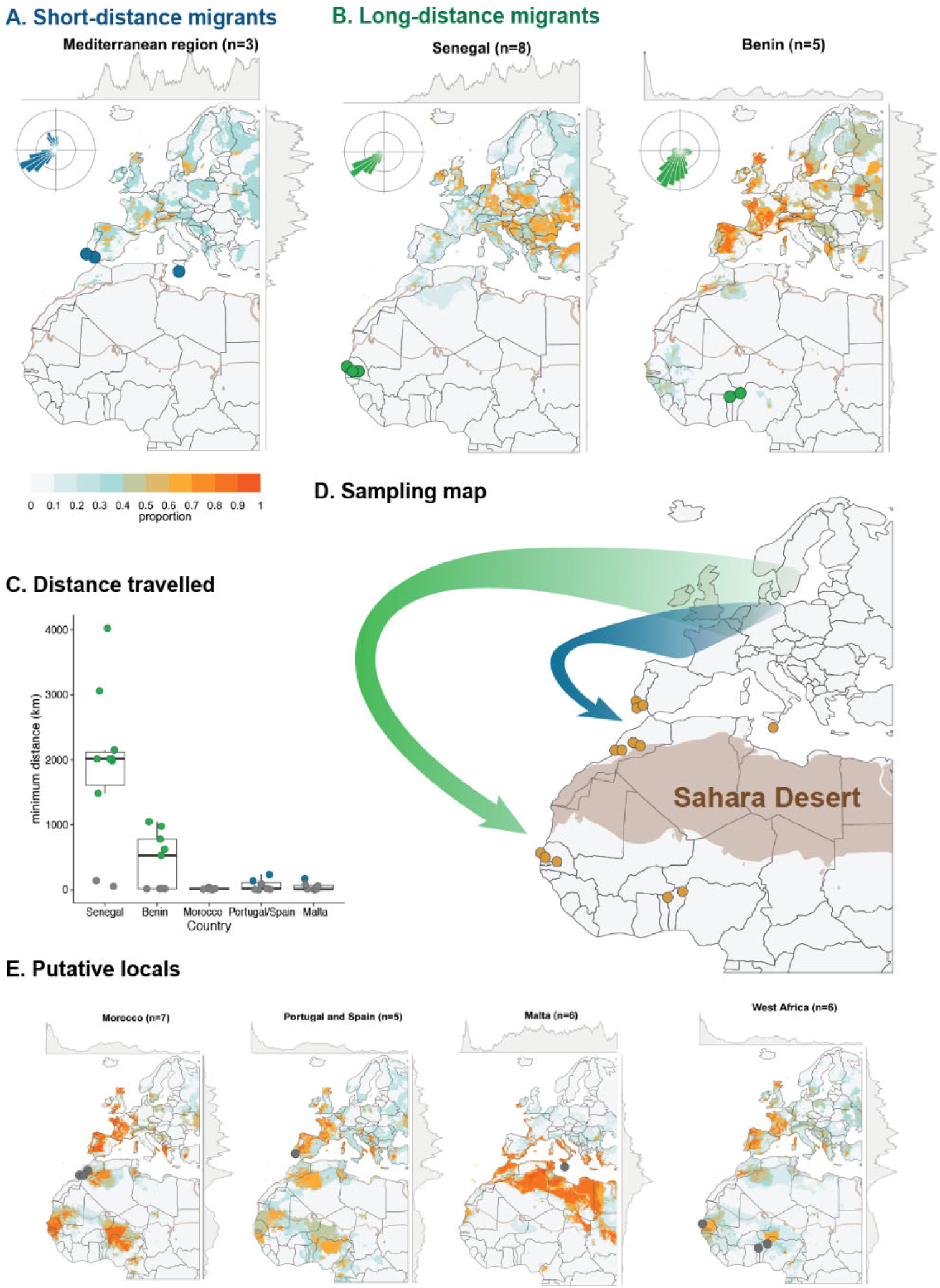
Summarised posterior probability surfaces and subsequent estimates of migration distance (i.e., minimum distance) and direction based on dual δ²H and ^87^Sr/^86^Sr-based geographic assignment. **(A,B,E)** Summary maps depict the proportion of individuals with a high probability of natal origin in a given area, as defined by the 2:1 odds ratio. The side margins display the mean proportion by latitude and longitude and only one circle per capture location is shown. The inset rose plot illustrates the average probability-weighted bearing from the estimated natal origin (centre of plot) to the capture locations. Maps summarise the results for **(A)** short-distance migrants, **(B)** long-distance migrants, and **(E)** putative locals captured in the circum-Mediterranean region and West Africa. **(C)** Boxplot displaying the estimated migration distance (km) categorised by the country of capture. Individuals with a minimum distance of less than 100 km were considered putative locals. **(D)** Map indicating the capture locations both north (Morocco, Portugal and Spain, and Malta) and south (Senegal and Benin) of the Sahara. Butterflies captured south of the Sahara generally migrated longer distances (green arrow) compared to those captured north of the Sahara (blue arrow).

Based on the estimates of migration distance, all individuals were grouped into three categories: (i) long-distance migrants (n = 13, 33%; with minimum migration distances ranging from over 500 to over 4,000 km), (ii) short-distance migrants (n = 3, 7%; with minimum migration distances ranging from over 140 to over 240 km), and (iii) individuals of putatively local origin (n = 24, 60%; having potentially travelled less than 100 km). Isotopic signatures corresponding to long-distance migrations were found exclusively in individuals captured south of the Sahara, specifically in Senegal and Benin. Based on the summarised posterior probability surfaces, these individuals seemed to originate either from northwestern Africa or Europe, notably from Portugal, Spain, France, or Sweden (Fig 1B). However, some of the individuals (n = 6, representing 32% of the samples from south of the Sahara; Fig 1E) had a high probability of originating from areas near their capture points (< 100 km) and were designated as “putative locals.” Notably, no individuals from south of the Sahara were classified in the short-distance migration category.

Among the samples collected north of the Sahara, some belonged to the short-distance migration category (n = 3, 14%; Fig 1A). However, the majority of these samples exhibited isotopic signatures consistent with local origin (n = 18, 86%; Fig 1E). The isotope-based analysis doesn’t clarify whether this local signature indicates (i) pre-migratory, locally-sourced offspring of early immigrants to the region or (ii) immigrants from a distant region with an isotopic signature similar to their capture location. We also considered natural history observations to interpret these patterns. The majority of individuals collected in the circum-Mediterranean region during October and November are likely the descendants of earlier arrivals to the collection area. This inference is substantiated by the favourable breeding conditions in the region during late September and October (9,11), as well as the observed presence of eggs, larvae, and recently-emerged adults (i.e., expelling meconium) at the time of collection in Morocco, Malta, and Portugal. Thus, we categorise any samples with a high probability of originating near their capture location as putative locals.

To assess the genomic differentiation between migratory phenotypes, it is crucial to integrate the isotopic signature data with spatiotemporal information from individual samples. Consequently, for subsequent analysis, we grouped individuals that travelled long distances from Senegal and Benin, explicitly excluding individuals with putatively local origins, into the “long-distance” group. To balance sample sizes, we grouped individuals from north of the Sahara, encompassing both the short-distance (n = 3) and putatively local categories (n = 18), into the “short-distance” group, operating under the assumption that these putative locals are the next-generation offspring of short-distance migrants.

### Wing morphology and wear are uncorrelated with migration distance

Wing wear scores, a metric scaled from 1 (fresh) to 5 (extremely worn), are often used as a proxy for butterfly age and migratory status (8,31,32). In our study, we preferentially sampled butterflies with worn wings (i.e., higher wing wear scores) as it is assumed that high wing wear scores correlate to post-migratory status; thus, wing wear scores ranged from 2 (few scales lost) to 5 (extremely worn). These wing wear scores showed no correlation with migration distance (Spearman rank correlation, rho = -0.06, p = 0.70).

Butterfly wing size and shape are sometimes correlated with migratory ability (e.g., 33). Here, wing shape was uncorrelated with migration distance (p = 0.2) or the country where the sample was collected (p = 0.1). There was a near-significant relationship between wing shape and wing size (R^2^ = 0.30; F_1,27_ = 1.8, p = 0.07). Wing shape differed between the sexes (F_1,27_ = 3.4, p = 0.0002), with the wings of females being more disparate in shape than those of males (S2 Fig; pairwise comparison, p = 0.03). Despite the differences in wing shape between males and females, we did not detect any difference in migration distance by sex (Wilcoxon test; W = 127, p = 0.47). Additionally, wing shape was influenced by the date of collection, with earlier captures having smaller and more elongated wings (S2 Fig; F_1,27_ = 3.0, p = 0.001).

### Lack of population structure across the Sahara

We explored if the distinct differences in migration distances, coupled with the evident spatiotemporal separation of short- and long-distance migrants, might be reflected in genetic differences between them, which in turn can be an indicator of population structure driven by neutral processes or as early signs of adaptive divergence among migratory phenotypes. We assessed these patterns at the whole-genome level, employing a comprehensive approach that integrated summary statistics, principal component analysis (PCA), and admixture proportion analysis. Remarkably, all approaches yielded consistent results: a lack of detectable population structure within the dataset and negligible differentiation between groups classified as short- and long-distance migrants.

First, using 13,215,545 high-quality callable sites from both coding and non-coding regions (38 individuals, two removed due to quality issues), we summarised levels of nucleotide diversity (π) within groups and genetic divergence (d_XY_) between groups of *V. cardui* individuals with distinct migratory behaviour (short-distance migrants: n = 21; long-distance migrants: n = 13), and calculated the fixation index (F_st_) as a measure of differentiation (S2 Table). We observed relatively high mean values of nucleotide diversity in both short- and long-distance migrants (π_autosome_ = 0.012 ± 0.003(1 SD)) compared to other Lepidoptera (34). Levels of the pairwise nucleotide divergence (d_XYautosome_ = 0.012 ± 0.003) between two phenotypes were elevated, reflecting the effect of high levels of diversity within samples and stochasticity of the estimates. The level of genetic differentiation between autosomes was low, but significant (pairwise F_ST_ = 0.001 ± 0.006, p < 0.001). A slightly higher fixation index was observed for the Z-chromosome (pairwise F_ST_ = 0.009 ± 0.016), which is expected given that the smaller effective population size makes the Z-chromosome more susceptible to the effects of genetic drift. We repeated this analysis using groups based on capture location (i.e., north vs. south of the Sahara), but found no substantial differences between summary statistics (S2 Table). These findings indicate that *V. cardui* that migrate short-versus long-distances have limited genetic differences and share similar distribution of allele frequencies, and that the effective population size in general is large.

Secondly, we used PCA to investigate potential genetic structuring. After additional filtering steps, we narrowed down our dataset to 813,810 high-quality single nucleotide polymorphisms (SNPs). This analysis revealed that the majority of samples clustered together with no separation between short- and long-distance migrants (Fig 2A), suggesting an overall absence of population structure. However, within this general pattern, two individuals stood out as outliers in the PCA; one showed differentiation primarily along the first principal component (i.e., 19H128) and the other exhibited differentiation along the second principal component (i.e., 19H115). Both of these outlier samples were collected in September and were among the earliest arrivals to sub-Saharan Africa (9; S1 Table). Moreover, these individuals had undertaken particularly long migratory journeys, with one covering over 2,000 km across the Sahara from northwest Africa or Europe (19H115) and the other over 1,000 km from Portugal, France, Sweden, or Finland (19H128) (S3 Fig). The clear separation of these individuals in the PCA biplot could be indicative of population structuring. We explored the contribution of specific SNPs to the first two principal components with *pcadapt* and identified 924 outlier SNPs that made significant contributions to the observed segregation of individuals. However, it is essential to note that drawing conclusions about the potential processes underlying the observed differences based on data from just two individuals is inherently limited. Nevertheless, we have provided a list of candidate SNPs that may serve as a valuable starting point for future investigations (see Data Availability Statement).

**Fig 2.**
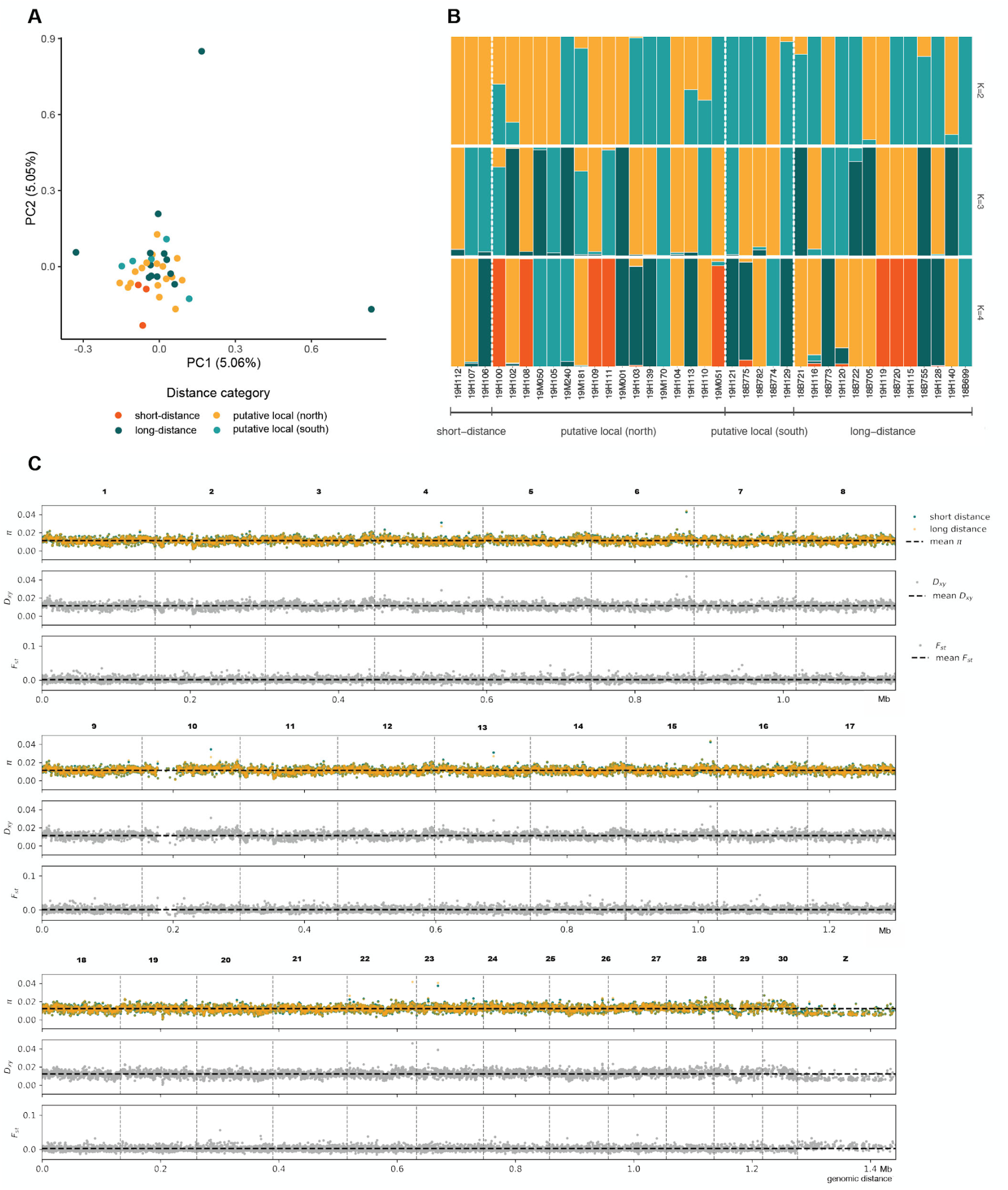
Lack of population structure and genetic differentiation between short- and long-distance migrants. **(A)** Principal component analysis (PCA) biplot illustrating the genetic variation in the samples; **(B)** Genomic admixture patterns based on assumptions of 2 to 4 ancestral populations (K = 2 to 4). No discernible structure was observed, with the optimal model demonstrating K = 1 (not visually depicted); **(C)** Genome scans showing within-population diversity (π), genetic differentiation (d_XY_) and fixation index (F_ST_) between individuals categorised as short- and long-distance migrants. The analysis was performed with sliding, non-overlapping windows of 20 kb. Mean values are indicated by a black dashed line.

Finally, we tested for population structure in our dataset using two different software implementations to infer individual ancestry proportions (admixture analysis). Specifically, we employed the sparse non-negative matrix factorization clustering algorithm (SNMF) and fastSTRUCTURE to test models assuming 1 to 5 subpopulations. In both cases, the best model was K = 1, indicating that the data are best described by a single panmictic population (Fig 2B).

### No islands of genetic differentiation between short- and long-distance migrants

The assessment of genetic structure using genome-wide averages may obscure regional differences in genetic differentiation. We therefore used window-based analysis of genetic differentiation between populations (i.e., F_ST_ scans) to quantify genetic differentiation across the genome and search for potential signatures of selection. Our analyses were based on two sample groupings: (i) migratory phenotype (i.e., short- and long-distance migrants) and (ii) capture location (i.e., north and south of the Sahara). Both comparisons revealed uniform differentiation landscapes characterized by a low average pairwise F_ST_. For autosomal regions, the average F_ST_ was 0.001 and regional peaks had a maximum F_ST_ of 0.113 (Fig 2C; S4 Fig). For the Z-chromosome, the average pairwise F_ST_ was 0.009 with a maximum peak of 0.072. This level of variation is consistent with stochasticity of the normalisation and/or neutral variation, as has previously been demonstrated in simulation studies (35) following simulation regimes applicable to our system. Additionally, none of the local F_ST_ peaks were accompanied by a simultaneous elevation of the genetic divergence and drop in diversity within each population extended over a substantial region, therefore not meeting the expectations for signatures of divergent selection.

## Discussion

### Contemporary evidence of variation in migratory behaviour

We used isotope-based estimates of migration distance to quantify the natural variation in migration distance exhibited by *V. cardui.* Our findings reveal that butterflies captured in Senegal and Benin during the autumn likely embarked on long migratory journeys, covering up to over 4,000 km as they crossed the Mediterranean Sea and Sahara from Europe. In contrast, *V. cardui* individuals captured in the circum-Mediterranean region in the same season had either undertaken shorter migrations or had isotopic signatures compatible with their capture area (Fig 1). Despite the increasing evidence that *V. cardui* was involved in regular trans-Saharan crossings as part of its annual migratory cycle, this evidence was limited to the monitoring of migratory arrivals in the sub-Sahara (9,10) and δ²H-based geolocation (8,36). We have expanded this isotope-based evidence with the addition of ^87^Sr/^86^Sr, narrowing down the potential geographical sources of these long-distance migrants and confirming that distinctive migration strategies are displayed by *V. cardui* found north and south of the Sahara during the autumn months.

In our study, we found that estimates of migration distance cannot be explained by wing size, wing shape, or sex. In this regard, *V. cardui* differs from North American monarch butterflies *Danaus plexippus*, which show strong associations between migratory ability and wing morphology in that larger and elongated wings are associated with the increased flight potential of the “super generation” during autumn migration (33,37,38). The variation in migration distance detected in our study appears to have a basis in behaviour rather than wing morphology, which may indicate limited selection on migration distance, as behaviour is thought to be more plastic than morphology (39).

We additionally found no significant association between wing wear score and migration distance. Although wing wear scores have long been used as a proxy for butterfly age and migratory status, with high wing wear scores corresponding to older butterflies that have completed their migratory journeys (e.g., 26,31,40), our results further illustrate that this proxy is highly unreliable (37). For example, the butterfly with the highest estimated migration distance, up to over 4,000 km (i.e., 18B721), had a low wing wear score of 2 (i.e., “few scales lost”; 32), which is often interpreted as belonging to a recently-emerged adult. This finding is in agreement with Korkmaz et al. (41), who conjecture that high altitude, gliding flight during migration results in less wing damage than low-altitude foraging and breeding behaviour.

### Trans-Saharan panmixia

Our genome-wide analysis revealed no evidence of genetic divergence between *V. cardui* individuals migrating to north or south of the Sahara during autumn (Fig 2). Consequently, the *V. cardui* population stretching from Northern Europe to the equatorial regions of sub-Saharan Africa can be characterised as regionally panmictic, confirming the spatial extent of the Afro-Palearctic migratory range. Our findings align well with previous studies using mitochondrial DNA markers (42–44). Given that large biogeographic barriers, such as the Mediterranean Sea and Sahara, do not seem to lead to population structure, it is likely that panmixia will be found at even larger scales. However, *V. cardui* migration seems to generally follow latitudinal trajectories, and extending to larger geographical scales might reveal longitudinal structure and population differentiation (e.g., 45). Future studies will need to assess the extent to which global panmixia occurs across the entire distributional range of *V. cardui*, preferably by including genome-wide genetic markers from around the world, including remote islands which sometimes host isolated populations that are phenotypically and genetically differentiated (e.g., 46).

Special attention should be given to understanding how selection and local adaptation act on migratory behaviour in insects, as well as the methods for detecting them. In our study, although we did not detect clear signatures of selection or adaptation, we did detect two outlier samples in the PCA. There could be multiple explanations for the patterns we detected. First, unlike migratory vertebrates that complete round-trip migrations within a single generation, insect migratory cycles are multi-generational. Multi-generational migratory insects exhibit a complex reticular movement pattern driven by the drastic changes in the location and size of the suitable breeding habitat over the annual migratory cycle (9,11). Therefore, polygenic within-generation selection due to spatially- and temporally-varying selection pressures could lead to allele frequency changes for a portion of the population (47). However, the same alleles selected for in one generation are unlikely to be favoured by selection pressures in the subsequent generation because of spatiotemporal differences in abiotic and biotic conditions during the migratory journey and in the novel breeding habitat, thus diluting the effect of within-generation local selection. Similar frameworks have been proposed for some fish species which show signs of within-generation local selection within panmictic populations (47–49). A similar scenario has been outlined for *D. plexippus*, which experiences selection pressures on wing shape and wing size during long-distance autumn migrations, only to have these pressures released in subsequent generations (37).

Second, it is important to recognize that the exceptionally large effective population size of *V. cardui* (50) and, consequently, weak genetic drift could impede our ability to detect early, or complete, reproductive isolation between short- and long-distance migrants. In these circumstances, many thousands of generations of reproductive isolation would need to elapse before our analysis would be able to detect significant allele frequency shifts. Therefore, it is possible that reproductive isolation has indeed occurred, but remains undetected. Third, it is conceivable that our sampling strategy was unable to detect genetic differentiation between groups of migrants. Notably, the most genetically differentiated migrants (Fig 2A) were on the leading edge of arrivals to West Africa (September 2019) (9) and likely originated in temperate Europe (S3 Fig). These early arrivals may have been exposed to strong environmental cues in temperate Europe, where seasonal changes occur earlier than in regions closer to the equator, prompting the butterflies to initiate their migrations earlier in the season. Early migrants of *D. plexippus* have been shown to have advantages over their later counterparts, such as enhanced immune defences (51). Consequently, it is possible that *V. cardui* segregates temporally, rather than by distance. However, it is more plausible that this temporal pattern results from within-generation selection, which is likely to be diluted in the subsequent generation. Despite any ephemeral within-generation local selection, we anticipate that the capacity to tolerate heterogeneous environments relies primarily on phenotypic plasticity (52). Ultimately, interpreting our results as indicative of regional panmixia remains the most parsimonious approach.

Panmixia across an Afro-Palearctic distributional range and regular trans-Saharan crossings are exceptional phenomena for insects, as evidenced by the fact that the Palearctic and Afrotropical regions are distinct biogeographic realms with unique faunal compositions, sharing only a relatively small number of species. We found that neither differences in migration distance nor the potential biogeographic barrier led to isolation, selection, or significant allele frequency shifts in any genomic region (Fig 2C). This finding is in contrast to numerous examples in vertebrate species (e.g., 15,17,53,54) and a few instances in insect species (e.g., 55). Some of the species of insects shared between the Palearctic and the Afrotropical regions are suspected to be migratory, particularly among the Lepidoptera, Odonata, and Orthoptera (56,57), and thus it is likely that trans-Saharan gene flow occurs in some of these other species as well. Continental-scale panmixia has been demonstrated in other long-distance migratory insect species, suggesting that panmixia is a prevalent feature among migratory species (but see 58). For example, whole-genome resequencing of *D. plexippus* revealed high gene flow and low genetic differentiation between butterflies located east and west of the Rocky Mountains, despite following geographically separated migration routes (59). Regional panmixia has also been suggested through microsatellite-based population genetics for the green darner dragonfly *Anax junius* in eastern North America (60,61) and global panmixia has been inferred through mitochondrial DNA analysis for the wandering glider dragonfly *Pantala flavescens* (62,63).

The widespread panmixia observed in many migratory insects raises questions about the life history strategies promoting panmixia. First, most insect migrations are multi-generational, featuring complex movement patterns driven by dynamic changes in breeding habitat, promoting intra-population mixing. Additionally, the tendency of long-distance migratory insects to engage in multiple matings, their lack of courtship behaviour, and their practice of laying numerous eggs (e.g., 60,64,65) hints at an eponymous, yet underexplored, link between panmictic tendencies and mating strategies in migratory insects. Large census population sizes in migratory insects, accompanied by intermittent outbreaks, further drive them towards panmixia. Outbreaks increase the number of successful migrants, potentially eroding local adaptation (66). Outbreaks of *V. cardui* from regions with anomalously high vegetation growth (67) may contribute to the observed lack of genetic structure. Similar population dynamics observed in species like *Libytheana carinenta* (68), *A. junius* (69), and some hoverflies (70) hint at outbreaks being pivotal in shaping the population structure of many migratory insects.

The same life-history factors of migratory insects that appear to promote panmixia among migratory insects (i.e., multi-generational migrations, an absence of assortative mating, and large census population sizes with outbreak dynamics) are also associated with large effective population sizes and high genetic diversity. The multi-generational annual migratory cycle results in a greater number of mutations occurring within the population per unit time, compared to species with longer generation times, and large census population sizes are less susceptible to genetic drift, both of which encourage genetic diversity (71). Indeed, our samples showed relatively high genetic diversity (π_autosome_ = 0.012) compared to other butterfly species (34). This observation aligns with the general trend that migratory butterflies tend to possess higher levels of heterozygosity than sedentary species (50). Consequently, this abundant genetic variation provides opportunities for at least some individuals to acclimatise to the heterogeneous environmental conditions that are encountered during their multi-generational migrations, thereby facilitating survival.

Due to the obligate, multi-generational migrations with low philopatry, random mating, and large, outbreaking populations, local adaptation is implausible within the Afro-Palearctic population of *V. cardui*. We suggest that rather than adapting to short or long-distance migration, a plastic response to the environment determines the differences we observe in migratory phenotype. It is also possible that the differences in migration distance observed here could also be explained, in whole or in part, by extrinsic mechanisms, including differences in parasite load (72), energy storage and metabolism (73), wind-assistance and weather events (74), or even personality (75). However, studies on *D. plexippus* suggest that while migratory state (i.e., resident vs migratory) is genetically determined (76), variation in migration distance is more likely to be due to many loci of small effect or differential gene expression (59), likely controlled by epigenetic mechanisms (4,18,77). Variation in migratory behaviour is likely a response to multiple environmental cues such as resource availability, photoperiod, temperature, humidity, and host plant quality. Recent experiments have demonstrated differences in *V. cardui* gene expression, methylation, and gene regulation due to the density of larval conspecifics, larval food availability, and host plant availability for oviposition (78,79), suggesting a strong response to environmental cues. The transcriptional basis of differences in migration distance has been explored in the moth *Helicoverpa armigera* and in *D. plexippus*, yielding promising candidate genes (59,80). Future controlled laboratory studies with *V. cardui* are needed to disentangle the environmental triggers and molecular pathways of phenotypic plasticity in migration distance.

### Conclusion

Here, the combination of isotope-based phenotyping of migration distance and comprehensive whole-genome resequencing provided a unique opportunity to investigate phenotype-genotype associations of migratory behaviour in insects, a challenging endeavour until now. We found distinct differences in migration distance, with most individuals captured in Benin and Senegal exhibiting long-distance migrations of up to 4,000 km, traversing the Mediterranean Sea and Sahara from temperate Europe. In contrast, *V. cardui* captured in the circum-Mediterranean region at the same time either undertook short-distance migrations or displayed isotopic signatures indicative of local origins. These differences in migratory behaviour did not have a clear association with wing morphology or wing wear. Genome-wide comparisons between these migratory groups found negligible genetic differentiation, leading us to conclude that the Afro-Palearctic population of *V. cardui* is panmictic, with regular bidirectional trans-Saharan crossings during the annual migratory cycle. Given the life history traits of *V. cardui*, namely multi-generational migration, their mating strategy, and large population sizes, local adaptation appears implausible within this population. Consequently, future research should explore the role of environmental cues and differential gene expression in shaping the diversity in migration distance.

## Materials and Methods

### 1. Sample collection

Wild, adult *V. cardui* were captured from six countries located both north and south of the Sahara (S1 Table). South of the Sahara, butterflies were captured in Senegal and Benin from August to November of 2018 and 2019 (n = 19). North of the Sahara, butterflies were captured from Morocco, Malta, Portugal, and Spain in November 2019 (n = 21). We used behavioural, demographic and timing evidence to maximise the probability of targeting migratory individuals at these locations. First, we preferentially sampled butterflies with worn wings (i.e., higher wing wear scores) and showing reproductive behaviour (i.e., hilltopping or oviposition) because *V. cardui* typically delay their reproductive behaviour until migration has been completed (81). Second, demographic data in some of the sampling locations supported the arrival of migrants. A monitoring station in Senegal saw a rising number of *V. cardui* adults in the autumn (August through October) during 2018 and 2019, followed, but not preceded, by a peak in larval counts, suggesting that the sudden spike in adults in the autumn is a result of immigration to the region (9). Similarly, a monitoring station in Benin saw high numbers of adults in September and October, followed, but not preceded, by high larval counts in October and November (9). Besides this demographic evidence, the time of the year when these butterflies were collected matches the first estimated arrival of migrants at the sampling locations. South of the Sahara, monitoring data does not indicate the presence of *V. cardui* year-round (9). North of the Sahara, much of the circum-Mediterranean region is unsuitable for *V. cardui* development in July, August, and September (Morocco) due to high temperatures and low host plant availability (40), suggesting that these individuals represent the first arrival of migrants in the region (9, 40).

### 2. Isotopic analysis

#### 2.1 Hydrogen isotope values

Prior to δ²H analysis, a forewing from each *V. cardui* was soaked, with agitation, in three successive baths (1 h, 30 min., 10 min.) of 2:1 chloroform:methanol solution to remove surficial dust and lipids, which are known to introduce error into δ²H measurements (82,83). Samples were cut from the same membranous region of the forewing to reduce intra-individual variation from pigmentation (82) and wing veins (24). Samples were then weighed (0.162 ± 0.009 mg) into silver capsules and surface moisture was dried in a 40 °C oven. Non-exchangeable δ²H values were determined at the Ján Veizer Stable Isotope Laboratory (University of Ottawa, ON, Canada) using a comparative equilibrium approach (84). All measurements were taken using high temperature (1400 °C) flash pyrolysis (TCEA, Thermo Finnigan, Germany) with a helium carrier passed through a chromium-filled reactor and, after separation, introduced to a Delta V Plus IRMS (Thermo Finnigan, Germany) via a Conflow IV interface (Thermo Finnigan, Germany). A three-point calibration was used to calibrate the sample δ²H values: CBS (caribou hoof; -157 ± 0.9‰; 85), KHS (kudu horn; -35.3 ± 1.1‰; 85), and USGS43 (human hair; -44.4 ± 2.0‰; 86). Internal standards were measured to assess the quality of the measurements, including one keratin reference standard, USGS42 (human hair; measured: -75.3 ± 0.5‰, n = 4; standard: -72.9 ± 2.2‰ (86)), and two in-house chitin standards, ground and homogenised *Lymantria dispar* (measured: -64.4 ± 1.8‰, n = 6; long-term average: -64 ± 0.8‰) and Alfa Aesar chitin (measured: -22.8 ± 0.7‰, n = 4; long-term average: -22 ± 1.2‰). Based on within-run replicates of the internal standards and repeated sample measurements, precision is estimated to be about ± 2‰. All reported δ²H values are normalised to the Vienna Standard Mean Ocean Water - Standard Light Antarctic Precipitation (VSMOW-SLAP) standard scale.

#### 2.2 Strontium isotope ratios

To prepare for ^87^Sr/^86^Sr analysis, a single forewing from each butterfly was cleaned using pressurised nitrogen gas for 2 min. at 69 kPa to remove any surface contaminants (e.g., dust; 26). The wings were then digested in 1 mL of 16 M HNO_3_ (distilled TraceMetal^TM^ Grade; Fisher Chemical, Canada) using microwave digestion (Anton Paar Multiwave 7000; Austria). The samples were heated from ambient temperature to 250 °C at a steady rate over 20 min. and then left at 250 °C for 15 min. in a pressurised chamber. An aliquot of 200 µL from each sample was separated and diluted to 2 mL of 2% v/v HNO_3_ and analysed for Sr content via inductively coupled plasma mass spectrometry (ICP-MS; Agilent 8800 ICP-QQQ, Agilent Technologies Inc., CA, USA) at the Department of Earth and Environmental Science, University of Ottawa (ON, Canada). Calibration standards were prepared using single element certified standards purchased from SCP Science (Montreal, Canada). Samples had an average of 3.1 ± 1.7 ng Sr per mg of wing tissue (n = 40).

The remaining aliquot of sample digest was dried down and re-dissolved in 0.5 mL 6 M HNO_3_. The separation of Sr was processed in a 100 µL microcolumn loaded with Sr-spec Resin (100 - 150 µm; Eichrom Technologies, LLC). The matrix was rinsed out using 6 M HNO_3_, and Sr was collected with 0.05 M HNO_3_. To ensure complete separation, the microcolumn process was repeated. After separation, eluates were dried and re-dissolved in 200 µL 2% v/v HNO_3_ for ^87^Sr/^86^Sr analysis.

The ^87^Sr/^86^Sr analysis was performed at the Pacific Centre for Isotopic and Geochemical Research (University of British Columbia, BC, Canada) using a Nu-Plasma II high-resolution MC-ICP-MS (Nu Instruments) coupled to a desolvating nebulizer (Aridus II, CETAC Technologies). The isotopes ^84^Sr, ^86^Sr, and ^87^Sr have isobaric interferences from ^84^Kr, ^86^Kr, and ^87^Rb respectively, and were corrected for using the ^85^Rb and ^83^Kr signals. Instrumental mass fractionation was corrected by normalising ^86^Sr/^88^Sr to 0.1194 using the exponential law (87). Procedural blanks were negligible. The reproducibility of the ^87^Sr/^86^Sr measurement for 5 ppb NIST SRM987 is 0.71025 ± 0.00009 (1 SD, n = 138) and for 1.4 ppb NIST SRM987, 0.71019 ± 0.00011 (n = 48). A matrix-matched chitin internal standard, 5 ppb Alfa Aesar chitin, was also used (0.713959 ± 0.00009; n = 3).

#### 2.3 Geographic assignment

Continuous-surface isotope-based geographic assignment was performed via the assignR package (88) in R v4.1.0 (89). The origins of the unknown origin *V. cardui* were estimated through a probabilistic framework using previously-calibrated isoscapes: (i) a hydrogen isoscape calibrated with residential butterfly samples from across the Palearctic and Afrotropical regions (90), and (ii) a regional bioavailable strontium isoscape (91). The most likely natal origin of each individual was estimated using δ²H isotopes alone, ^87^Sr/^86^Sr isotopes alone, and using both isotopes together (see Data Availability Statement). The 2:1 odds ratio was chosen, in keeping with previous studies (e.g. 31), to create binary surfaces from the posterior probability surfaces of the dual δ²H and ^87^Sr/^86^Sr-based geographic assignment, representing the top third of the probability distribution as high probability of natal origin (i.e., “1”) and everything else as low probability of natal origin (i.e., “0”). Individual binary surfaces were then summed for each capture location, and divided by the number of individuals, to summarise the results of the geographic assignment into maps of the proportion of individuals with a high probability of natal origin at a given raster cell.

#### 2.4 Distance metric

A consensus has not been reached on the optimal method by which to estimate the distance a migratory individual has travelled from posterior probability surfaces (92). We assessed the ability of seven different metrics to accurately estimate the migration distance of aerial migrants in our study area using synthetic data; we generated isotopic signatures for 250 synthetic migrants, performed dual δ²H and ^87^Sr/^86^Sr geographic assignment for each synthetic migrant, and calculated each of seven distance metrics. Most metrics were highly inaccurate, and could under- or over-estimate the actual great circle distance by thousands of kilometres (S1 Fig). However, the “minimum distance” metric consistently underestimated migration distance for our study area and, therefore, acts as a conservative estimate of migration distance (S1 Fig). Therefore, in this study, we used the minimum distance metric to estimate the distance travelled. This metric measures the shortest distance between the capture location and an isopleth threshold, here defined as the top third of the probability distribution (0.3 isopleth; e.g., 93,94).

Many butterflies had an isotopic signature similar to the local isotopic signature of their capture location. We interpreted individuals with a highly probable natal origin at the capture location (as defined by the 2:1 odds ratio) and/or that had a minimum distance less than 100 km as “putative locals.” These individuals are likely either the locally-sourced offspring of the previous migratory generation or were immigrants from a far-off region with a similar isotopic signature to that of their capture location. Our interpretation of these samples as putative locals is the more parsimonious approach.

### 3. Geometric morphometric analysis

Differences in migration distance or trajectories could potentially be explained by intrinsic factors such as sex, body size, or wing shape. To test for differences in wing shape, photographs of each sample’s forewing were taken and used for morphometrics analysis. Since we preferentially chose samples that had higher wing wear scores, extensive damage to the wings made landmarking of many samples impossible. Therefore, only 28 of the 40 samples were landmarked using TPSDig2 v.2.32 (95) using 12 conventional landmarks for Nymphalid butterflies (e.g., 96,97). Using the R package geomorph (98), a generalised procrustes analysis removed the translational, rotational, and scaling variation of the shape. A procrustes regression was used to test the effects on wing shape of sex, month of collection, estimated migration distance, and butterfly wing size.

### 4. Genomic analysis and genotyping

#### 4.1 Whole genome resequencing

Library preparation for all 40 samples was done using TruSeq PCR-free kits (Illumina, San Diego, USA) at the Institut Botànic de Barcelona (IBB – CSIC, Barcelona, Spain). Whole-genome resequencing was performed with paired-end 2x151 bp reads on one Illumina NovaSeq 6000 S4 lane at the National Genomics Infrastructure (NGI; Stockholm, Sweden) to generate a mean depth of coverage per individual of about 48 reads.

#### 4.2 SNP calling

A modified Sarek Nextflow pipeline (99) was used for preprocessing of the whole genome resequencing data and for detecting SNPs. At the first step of the Sarek pipeline, we evaluated the quality of the raw reads with FastQC (version 0.11.9; 100). Reads were stringently trimmed using fastp (101,102) to remove (i) sequences from the 5’ end (12 bp) and 3’ end (3 bp) of each read, (ii) adaptor sequences, (iii) low quality reads (overall Phred score < 30), and (iv) short reads (< 30 bp). Read quality after trimming was re-evaluated using FastQC. At the next step, a mean of 96.6% of the reads were mapped to the reference genome (103) using BWA-MEM (version 0.7.17; 104) and then sorted and indexed using SAMtools (version 1.9; 105). Optical and PCR duplicates were marked using GATK MarkDuplicates (version 4.1.4.1; 106). These duplicates, as well as sequences with low mapping quality (MAPQ < 30), were removed using SAMtools. Since prior SNP data for *V. cardui* were unavailable, the base calibration (BQSR) step module in the Sarek pipeline was skipped.

The first steps of variant calling were performed using GATK Best Practices (107), as implemented through the Sarek pipeline. In order to decrease computational time, the Sarek pipeline calls each chromosome separately. MultiQC (version 1.8) was used to assess the quality of the output from all steps of the pipeline. The resulting per chromosome vcf files included from 1.3 to 3 million unfiltered SNPs (78.9 million SNPs total). The last step of variant calling was performed outside of the pipeline to accommodate for invariant site calling (-all-sites option in GenotypeGVCFs), which is necessary for reliable downstream estimation of population genetic summary statistics, such as π and d_XY_. This variant filtering included multiple steps: (i) hard filtering, (ii) removing SNPs overlapping with TE annotation, (iii) separate quality and coverage filtering for variant and invariant sites, and (iv) merging of variant and invariant site vcfs for the F_ST_ estimation. Hard filtering, which is meant to substitute for the variant quality score recalibration (VQSR) step in GATK Best Practices (107), was performed on raw vcf files, which included both variant and invariant sites. Using bcftools v1.15 (108), we filtered for variant quality by depth (QD < 2.0), Fisher strand bias (FS > 60.0), root mean square mapping quality (MQ < 40.0), u-based z-approximation from the Rank Sum Test for mapping qualities (MQRankSum < -12.5), and u-based z-approximation from the Rank Sum Test for site position (ReadPosRankSum < -8.0). We proceeded to remove indels and multivariate sites (--exclude-types indels, other --max- alleles 2) and reindexed the resulting vcf file. Next, the coordinates of transposable elements and simple repeats were obtained with a custom bash script from a previously published annotation of transposable elements (109), and used to remove SNPs appearing in these regions.

The quality of the resulting files (one vcf file per chromosome) was evaluated using summary statistics implemented with vcftools (i.e., allele frequency (--freq2), mean depth (-- depth), mean per site depth (--site-mean-depth), per site quality (--site-quality), per sample missingness (--missing-indv), per site missingness (--missing-indv), heterozygosity estimation (--het); 110). Two individuals were removed due to quality issues (Sample IDs: 19H137 and 19H114). Variant and invariant sites were split into two files for downstream analysis. Informed by the summary statistics, we applied filters for quality and depth to the variant sites vcf file (i.e., minor allele frequency (--max-maf 0), mean depth value (--min- meanDP 30), mean depth value (--max-meanDP 80)). We also applied additional filters to the invariant sites vcf file (i.e., quality value (--minQ 30), mean depth value (--min-meanDP 30)). Separate files with both variant and invariant calls were kept as separate chromosomes to decrease computational time. The W chromosome was also removed from the analysis. A region with divergent values of populational statistics was detected in the first third of chromosome 10 corresponding to a large area of transposable element proliferation (Fig 2C; 109). Since this region likely does not carry information about genetic structure it was also filtered from the population structure analyses. On a technical note, it is important to mention that this region was retained after stringent SNP filtration, and future research on *V. cardui* should be aware that this region may necessitate special treatment (e.g., manual filtering).

#### 4.3 Population genetic analysis

For PCA and admixture analysis, variant files per chromosome were concatenated. To investigate genetic variation between individuals, a PCA was performed using PLINK v1.90 b 4.9 (Fig 2A). For this analysis, we additionally filtered out singleton SNP sites (i.e., MAF = 0.03), and SNPs that were in linkage (i.e., SNPs with r^2^ values > 0.1 using sliding windows of 50 SNPs with a step size of 10 SNPs); a total of 813,810 high quality variants remained after filtering. Outliers in the first two principal components were detected using the R package pcadapt (111) using Bonferroni-corrected p-values (α = 0.05). The PCA was repeated excluding the two outliers, but no additional structure was detected (S5 Fig).

Reverse migrations from sub-Saharan Africa to the Maghreb are thought to occur in the autumn, even while southwards trans-Saharan migrations are still underway (8). All of the samples captured in Morocco were designated as putative locals, but the posterior probability surfaces of some showed a possibility of originating south of the Sahara (Fig 1E). To test whether these samples could have biased our results, we repeated the PCA without samples collected in Morocco (S5 Fig). The resulting PCA was based on 647,906 SNPs and showed a single cluster with the same outlier individuals as the original PCA.

After excluding SNPs on the Z chromosome and in coding sequences, genetic structure was also assessed with a likelihood method, the sparse non-negative matrix factorization clustering algorithm, SNMF, in the R package LEA v3.6.0 (112). Each number of population clusters (K) from 1 to 5 was run independently with 5 repetitions with the regularisation parameter set to 100, after which cross-validation errors calculated using the cross-entropy criterion were compared. This process was repeated using a Bayesian framework via fastSTRUCTURE (113) using the simple allele frequency prior with 5 repetitions. The number of population clusters from 1 to 5 that best explained the observed structure was chosen with the chooseK.py tool.

#### 4.4 Genomic scan for selection

To detect any fine-scale regions of differentiation along the genome, a genome scan was performed on groupings based on capture location (i.e., north vs. south of the Sahara) and migratory phenotype (i.e., long vs. short migration distance). For the grouping based on migratory phenotype, individuals from south of the Sahara that were putative locals (n = 6) were removed, while putative locals from north of the Sahara (n = 19) were included with the short-distance migrants (n = 3) to balance the sample size. For genome scans we retained SNPs falling into coding regions and having minor allele frequencies. We used pixy v1.2.7 (114,115) to calculate nucleotide diversity (π), absolute divergence (d_XY_), and genetic differentiation (F_ST_; Hudson method (-hudson)) in 20 kb non-overlapping windows using all sites. Windows with less than 10% of sites represented (i.e., < 2000 bp) were removed from the analysis. Resulting per window statistics were summarized using custom python scripts. For low levels of F_ST_, we tested if the index significantly differed from 0 using a t-test in the python scipy package (p-value: 1.3e-216).

## Acknowledgements

This study was funded by grant 2018-00738 of the New Frontiers in Research Fund (Government of Canada) to GT and CB; by grants PID2020-117739GA-I00 MCIN / AEI / 10.13039/501100011033 and WW1-300R-18 (National Geographic Society) to GT; and by grant LINKA20399 from the CSIC iLink program to NB, CB and GT. MR was supported by the Queen Elizabeth II Graduate Scholarship in Science and Technology (QEII-GSST), the Ontario Graduate Scholarship, and the 2021 ORIGIN project (NSF award DBI-1565128). NB is supported by a research grant from the Swedish Research Council FORMAS (grant 2019-00670). DS is supported by the Thurberg postdoctorate fellowship program of Swedish Collegium for Advanced Science (Knut and Alice Wallenberg Foundation). The authors acknowledge support from the National Genomics Infrastructure in Stockholm funded by Science for Life Laboratory, the Knut and Alice Wallenberg Foundation and the Swedish Research Council, and SNIC/Uppsala Multidisciplinary Center for Advanced Computational Science for assistance with massively parallel sequencing and access to the UPPMAX computational infrastructure. The computations were enabled by resources provided by the National Academic Infrastructure for Supercomputing in Sweden (NAISS) and the Swedish National Infrastructure for Computing (SNIC) in Uppsala, partially funded by the Swedish Research Council through grant agreement no. 2022-06725 and no. 2018-05973. We are grateful to members of the Backström lab and Sean Stankowski for insightful discussions about the genome scan analysis. We would like to thank Smita Mohanty for help with the ICP-MS, Kerry Klassen and Paul Middlestead at the Ján Veizer Stable Isotope Laboratory for help with the hydrogen isotope analysis, and Kathy Gordon at the Pacific Centre for Isotopic and Geochemical Research for help with the strontium isotope analysis. We would like to thank Leonardo Dapporto, Leonardo Platania, and Aurora García-Berro for sharing code and knowledge related to the wing geometric morphometrics analysis, and Eude Goudégnon, Roger Vila, Mattia Menchetti, and Tomasz Suchan for help in sample collection. We would also like to thank Andrea Contina, Gabriel Bowen, and Michael Wunder for sharing code and insight relating to the migration distance estimates.

## Data Availability Statement

The raw sequence data is deposited in the European Nucleotide Archive (ENA) with accession number PRJEB70298. All data and code pertaining to the isotopic, geometric morphometric, and genomic analyses, including bash, R, and python scripts, can be found on the Open Science Foundation (https://osf.io/dzh2g/?view_only=6dc9f9bd496f4b538b0708d5f13cfce8).

