## Supporting Information for "Isotope geolocation and population genomics in *Vanessa cardui:* Short- and long-distance migrants are genetically undifferentiated"

**S1 Table. Sample metadata.** Capture date and location of each *V. cardui* sample and the hydrogen isotope value ($\delta^2H$) and strontium isotope ratio ($^{87}Sr/^{86}Sr$) measured in the wing tissue.

| Sample | Capture Date | Latitude | Longitude | Country | $\delta^2H$ (± SD) | $^{87}Sr/^{86}Sr$ (± SD) |
|---|---|---|---|---|---|---|
| 18B755 | 22.IX.2018 | 11.70681 | 3.216345 | Benin | -81 ± 2 | 0.71318 ± 0.00003 |
| 18B782 | 22.IX.2018 | 11.70681 | 3.216345 | Benin | -79 ± 2 | 0.71367 ± 0.00006 |
| 18B773 | 23.IX.2018 | 11.74402 | 3.276339 | Benin | -84 ± 2 | 0.71298 ± 0.00009 |
| 18B774 | 23.IX.2018 | 11.74402 | 3.276339 | Benin | -81 ± 2 | 0.71516 ± 0.00004 |
| 18B775 | 23.IX.2018 | 11.74402 | 3.276339 | Benin | -76 ± 2 | 0.71327 ± 0.00005 |
| 19H129 | 15.VIII.2019 | 10.7275 | 1.380323 | Benin | -77 ± 2 | 0.72074 ± 0.00006 |
| 19H128 | 15.IX.2019 | 10.7275 | 1.380323 | Benin | -83 ± 2 | 0.72613 ± 0.00006 |
| 19H137 | 9.X.2019 | 10.7275 | 1.380323 | Benin | -83 ± 2 | 0.71756 ± 0.00004 |
| 19H140 | 7.XI.2019 | 10.7275 | 1.380323 | Benin | -84 ± 2 | 0.71525 ± 0.00005 |
| 19M240 | 6.XI.2019 | 30.33404 | -9.4085 | Morocco | -68 ± 2 | 0.71288 ± 0.00004 |
| 19M001 | 31.X.2019 | 30.4975 | -8.82024 | Morocco | -74 ± 2 | 0.71375 ± 0.00002 |
| 19M002 | 31.X.2019 | 30.4975 | -8.82024 | Morocco | -87 ± 2 | 0.71353 ± 0.00004 |
| 19M170 | 4.XI.2019 | 31.38014 | -5.99294 | Morocco | -64 ± 2 | 0.71362 ± 0.00003 |
| 19M181 | 4.XI.2019 | 31.38014 | -5.99294 | Morocco | -72 ± 2 | 0.71473 ± 0.00002 |
| 19M050 | 1.XI.2019 | 31.73267 | -7.2829 | Morocco | -75 ± 2 | 0.71655 ± 0.00003 |
| 19M051 | 1.XI.2019 | 31.73267 | -7.2829 | Morocco | -79 ± 2 | 0.71451 ± 0.00002 |
| 18B720 | 13.IX.2018 | 13.97696 | -14.5436 | Senegal | -100 ± 2 | 0.71438 ± 0.00009 |
| 18B721 | 13.IX.2018 | 13.97696 | -14.5436 | Senegal | -109 ± 2 | 0.71522 ± 0.00005 |
| 18B722 | 13.IX.2018 | 13.97696 | -14.5436 | Senegal | -69 ± 2 | 0.71543 ± 0.00003 |
| 18B699 | 08.IX.2018 | 14.35961 | -16.4946 | Senegal | -101 ± 2 | 0.71063 ± 0.00011 |
| 18B705 | 07.IX.2018 | 14.71873 | -17.1292 | Senegal | -86 ± 2 | 0.71021 ± 0.00002 |
| 19H115 | 29.IX.2019 | 14.70977 | -17.1248 | Senegal | -79 ± 2 | 0.71013 ± 0.00002 |
| 19H116 | 29.IX.2019 | 14.70977 | -17.1248 | Senegal | -81 ± 2 | 0.70971 ± 0.00002 |
| 19H119 | 26.X.2019 | 14.70977 | -17.1248 | Senegal | -91 ± 2 | 0.70980 ± 0.00001 |
| 19H120 | 26.X.2019 | 14.70977 | -17.1248 | Senegal | -72 ± 2 | 0.71020 ± 0.00001 |
| 19H121 | 9.XI.2019 | 14.70977 | -17.1248 | Senegal | -65 ± 2 | 0.70997 ± 0.00002 |
| 19H104 | 18.XI.2019 | 35.90683 | 14.39475 | Malta | -57 ± 2 | 0.70992 ± 0.00002 |

| | | | | | | |
|---|---|---|---|---|---|---|
| 19H100 | 18.XI.2019 | 35.93002 | 14.40412 | Malta | -57 ± 2 | 0.71090 ± 0.00002 |
| 19H139 | 18.XI.2019 | 35.93002 | 14.40412 | Malta | -73 ± 2 | 0.71022 ± 0.00002 |
| 19H102 | 20.XI.2019 | 35.94727 | 14.35173 | Malta | -59 ± 2 | 0.70997 ± 0.00002 |
| 19H103 | 20.XI.2019 | 35.94727 | 14.35173 | Malta | -57 ± 2 | 0.71007 ± 0.00003 |
| 19H105 | 22.XI.2019 | 36.06184 | 14.28611 | Malta | -49 ± 2 | 0.70954 ± 0.00001 |
| 19H106 | 22.XI.2019 | 36.06184 | 14.28611 | Malta | -77 ± 2 | 0.71151 ± 0.00004 |
| 19H108 | 24.XI.2019 | 37.09005 | -8.16606 | Portugal | -69 ± 2 | 0.71025 ± 0.00004 |
| 19H109 | 24.XI.2019 | 37.24111 | -8.35311 | Portugal | -70 ± 2 | 0.71397 ± 0.00004 |
| 19H110 | 25.XI.2019 | 37.26194 | -8.5447 | Portugal | -71 ± 2 | 0.71369 ± 0.00004 |
| 19H111 | 25.XI.2019 | 37.26194 | -8.5447 | Portugal | -85 ± 2 | 0.71439 ± 0.00003 |
| 19H112 | 27.XI.2019 | 37.26194 | -8.5447 | Portugal | -90 ± 2 | 0.71779 ± 0.00004 |
| 19H113 | 27.XI.2019 | 37.26194 | -8.5447 | Portugal | -57 ± 2 | 0.71817 ± 0.00002 |
| 19H107 | 23.XI.2019 | 37.20373 | -7.26035 | Spain | -87 ± 2 | 0.71279 ± 0.00003 |

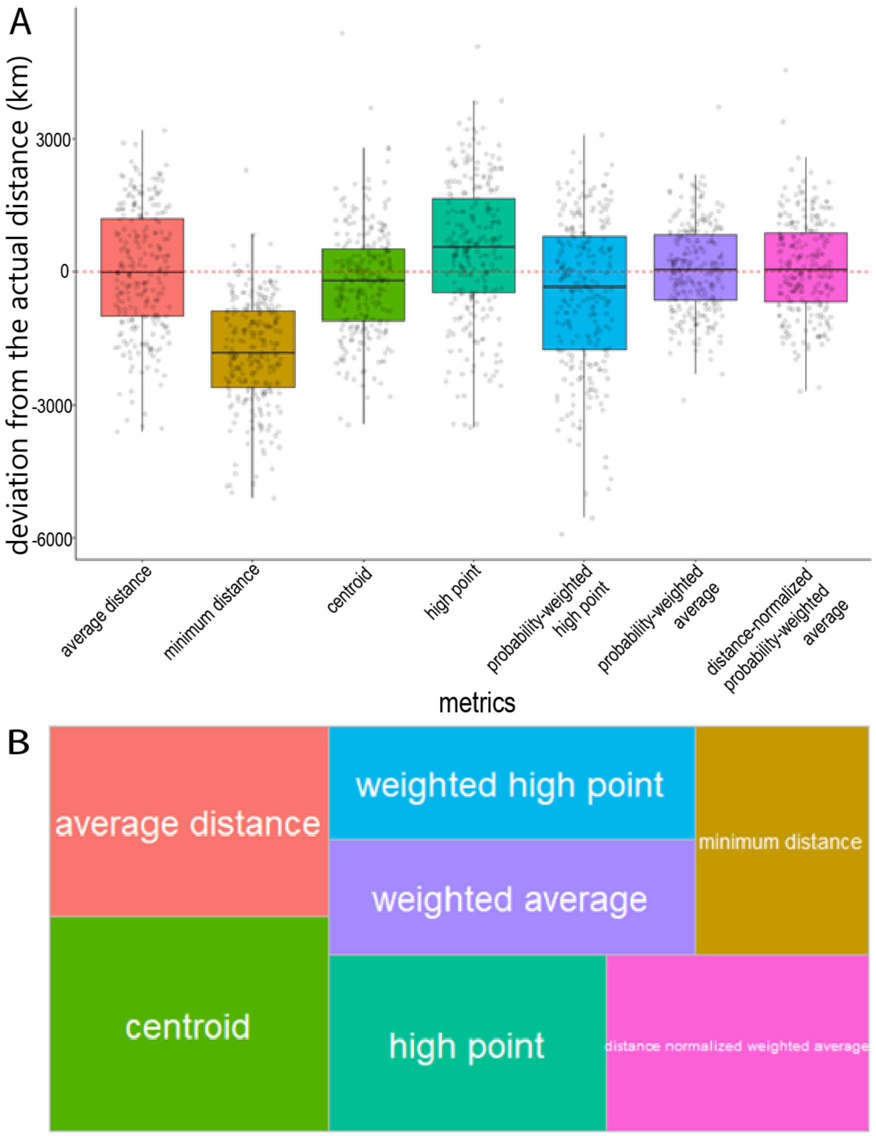


**S1 Fig. Performance of seven metrics estimating the distance travelled by 250 synthetic migrants.** Metrics include: (i) the average distance metric, which acts as the null model as it is determined mainly by geography, (ii) the minimum distance method, calculating the shortest distance to an isopleth threshold value (e.g., 94). (iii) the centroid method, calculating the distance to the centre of the highly probable area of natal origin, usually defined by the 2:1 odds ratio (e.g., 37), (iv) the high point method, calculating the distance to the raster cell with the highest probability (e.g., 92), (v) the weighted high point, calculating the probability-weighted distance to the raster cell with the highest probability, (vi) the probability-weighted average distance, calculating the probability-weighted average distance (assignR package: 88), and (vii) the distance-normalised probability-weighted average distance, calculating the probability-weighted average distance, while normalising for the number of cells at each distance. **(A)** Deviations from the actual distance (km) for each estimate of migration distance. Values below the red dashed line indicate under-estimates of the actual distance. **(B)** Treemap showing the proportion of synthetic migrants for which each metric was the best at estimating the actual migration distance. The average distance metric was the best at estimating migration distance for 16% of the individuals, demonstrating the inaccuracy of these metrics.

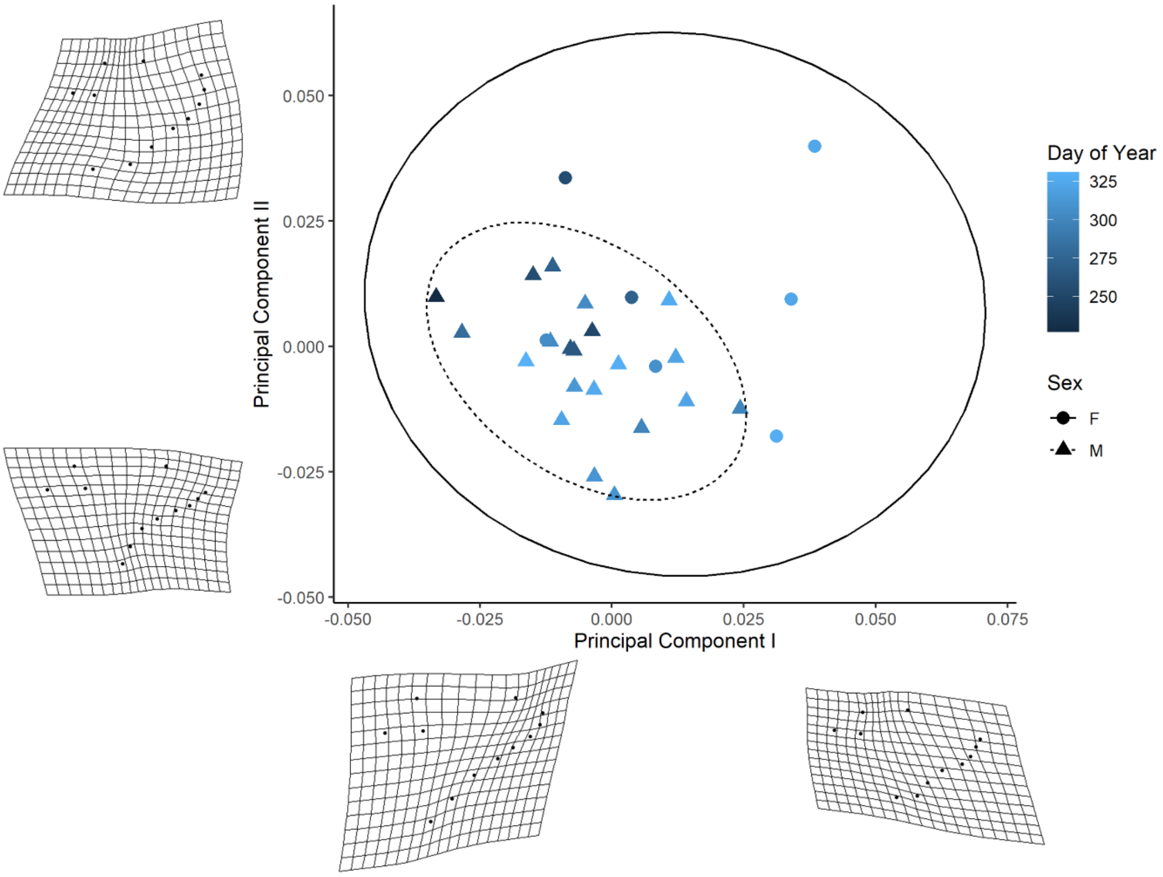


**S2 Fig. Wing geometric morphometric analysis.** Principal components biplot of *V. cardui* wing shape (n = 28). The shade indicates the day of the year that the sample was captured and shape indicates males (triangles) and females (circles). Females have greater variation in shape, as indicated by the larger 95% confidence ellipse for females (solid) compared to males (dashed). Shape differences between the extremes of the first principal component range from a subtly narrowed wing to a round wing (shape differences from the mean magnified by 5).

**S2 Table. Population genetic summary statistics.** Within-group nucleotide diversity ($\pi$) and between-group absolute divergence ($d_{XY}$) and genetic differentiation ($F_{ST}$) calculated separately for the autosomes and the Z chromosome (mean ± sd, maximum). Summary statistics are obtained from window-based values (20 kb) calculated in pixy. Samples are grouped by migration distance (i.e., short distance and Mediterranean putative locals vs long distance) and capture location (north vs south of the Sahara).

| | Migration distance | | Capture location | |
|---|---|---|---|---|
| | Autosomes | Z chromosome | Autosome | Z chromosome |
| $\pi$ (short/north) | 0.012 ± 0.003, < 0.070 | 0.008 ± 0.003, < 0.020 | 0.012 ± 0.003, < 0.070 | 0.008 ± 0.003, < 0.018 |
| $\pi$ (long/south) | 0.012 ± 0.003, < 0.067 | 0.008 ± 0.003, < 0.022 | 0.012 ± 0.003, < 0.067 | 0.008 ± 0.003, < 0.020 |
| $d_{XY}$ | 0.012 ± 0.003, < 0.069 | 0.008 ± 0.003, < 0.023 | 0.012 ± 0.003, < 0.069 | 0.008 ± 0.003, < 0.020 |
| $F_{ST}$ | 0.001± 0.006, < 0.113 | 0.009 ± 0.016, < 0.072 | 0.001 ± 0.005, < 0.048 | 0.007 ± 0.005, < 0.019 |

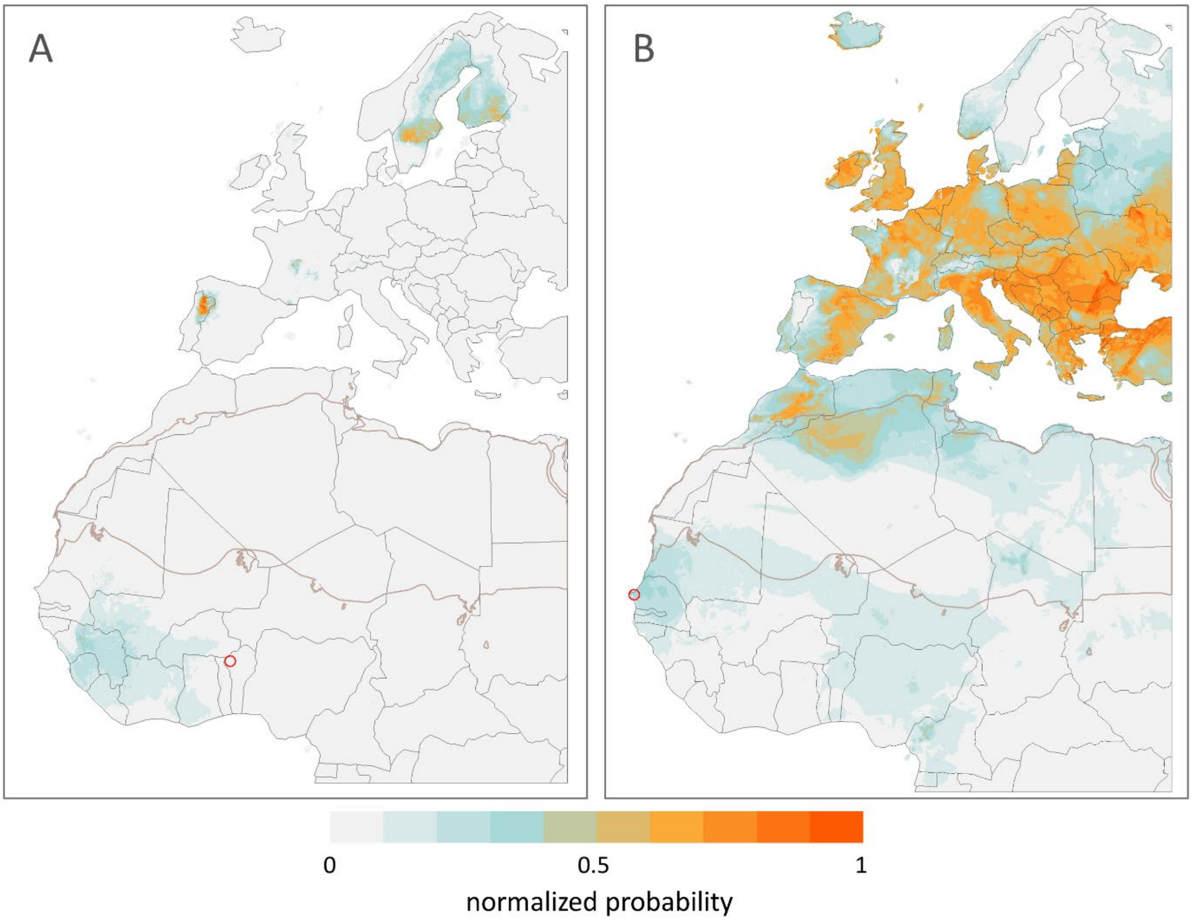


**S3 Fig. Examples of normalized posterior probability surfaces.** Samples **(A)** 19H128 and **(B)** 19H115 were collected at the red circles in September 2019.

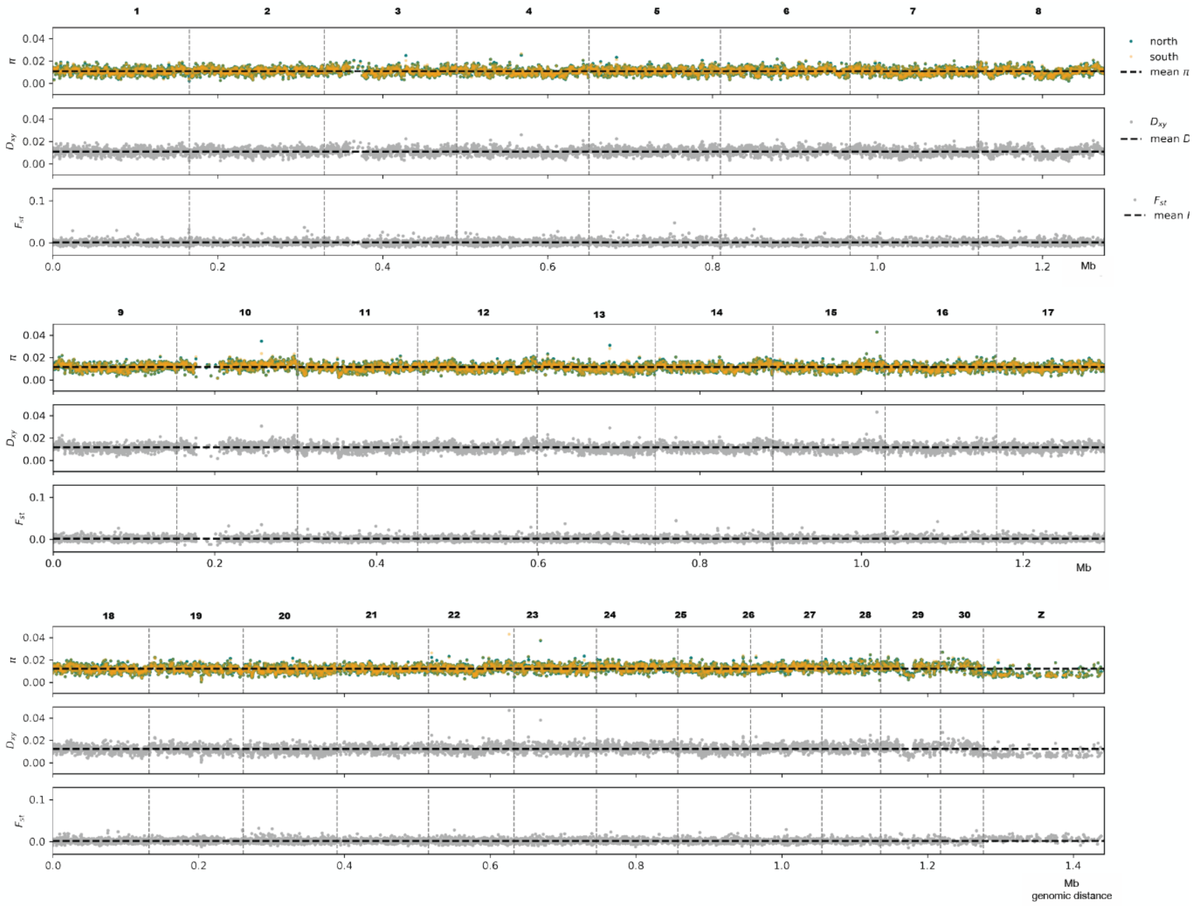


**S4 Fig. Genome scans showing within-population diversity (π), genetic differentiation ($d_{XY}$) and fixation index ($F_{ST}$) between individuals captured north and south of the Sahara.** The analysis was performed with sliding, non-overlapping windows of 20 kb. Mean values are indicated by a black dashed line.

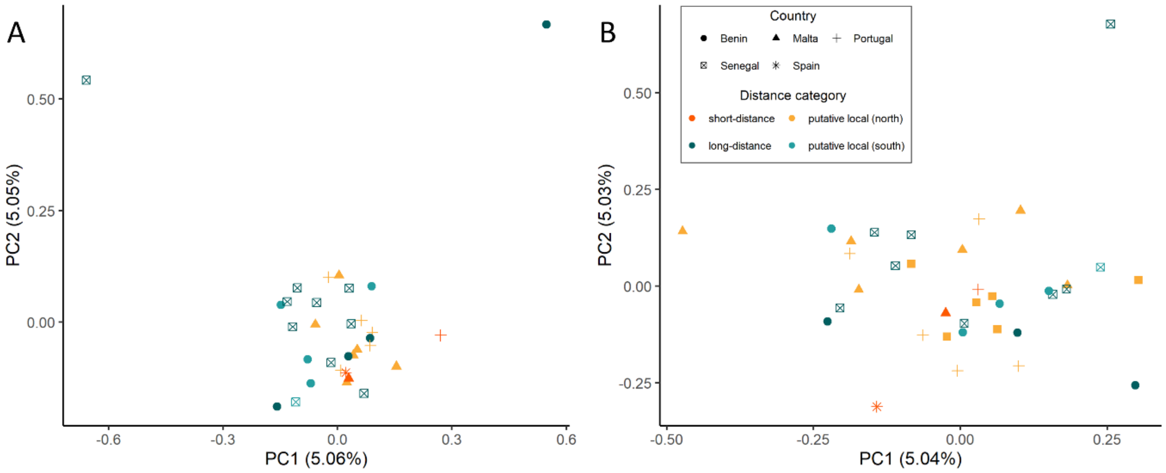


**S5 Fig. Principal component analysis (PCA) biplot illustrating the genetic variation in samples, (A)** excluding samples from Morocco, and **(B)** excluding two outliers (19H128 and 19H115).
